# In situ architecture of plasmodesmata suggests mechanisms controlling intercellular exchange

**DOI:** 10.1101/2025.07.24.666190

**Authors:** Marcel Dickmanns, Matthias Pöge, Peng Xu, Sven Gombos, Manuel Miras, Jürgen M. Plitzko, Rüdiger Simon, Waltraud X. Schulze, Wolf B. Frommer, Wolfgang Baumeister

## Abstract

Plasmodesmata are nanoscopic channels that traverse plant cell walls, establishing membrane and cytoplasmic continuity between adjacent cells to enable direct intercellular exchange. Although numerous components have been identified, their molecular organization and roles in controlling passage remain unclear. Here, we used cryo-electron tomography to resolve the in situ architecture of plasmodesmata in *Physcomitrium patens* across tissues and physiological states. We show how callose deposition at the cell wall shapes channel architecture to modulate permeability and identify helical protein assemblies formed by Multiple C2 domain and Transmembrane Proteins (MCTPs), which scaffold a central endoplasmic reticulum tubule and tether it to the plasma membrane. These findings define core architectural features of plasmodesmata and establish a structural framework for understanding how membrane, protein and cell wall components coordinate intercellular connectivity in plants.

## Introduction

Multicellular organisms rely on intercellular communication to coordinate growth, development, and responses to environmental cues (*1*). In plants, communication is complicated by the presence of a several hundred nanometers thick cell wall. To facilitate direct molecular exchange across this barrier, plants have evolved plasmodesmata (PD), nanoscopic intercellular channels with complex architectures. PD traverse the cell wall and establish continuity between the cytoplasm, plasma membrane, and endoplasmic reticulum (ER) of adjacent cells (*2*). In animal cells, direct cytoplasmic exchange largely occurs through transient clusters of protein channels known as gap junctions. Cytoplasmic bridges containing shared membranes, however, are rare and typically restricted to the germline such as in the ring canals of insect germ cell cysts (*1, 3*). In contrast, PD are ubiquitous across both germline and somatic plant tissues, forming an almost organism-wide continuum of cells. This continuity necessitates control of PD permeability to maintain cellular homeostasis and tissue identity (*4, 5*). Although PD are central to plant physiology and development, their molecular architecture and mechanisms governing permeability control remain incompletely understood. Proteomics-based approaches have identified hundreds of putative PD-associated proteins (*6*–*13*), but an integrated molecular model is still lacking. Genetic dissection remains challenging, as loss-of-function mutants often exhibit severe systemic phenotypes or lethality (*9, 14, 15*). In some cases, conditional expression systems or partial knockdowns have enabled functional studies, but genetic redundancy among PD-associated proteins further complicates analysis (*16*–*18*). Fluorescence microscopy allows tracking of candidate protein localization and dynamics but lacks the spatial precision to resolve protein organization at plasmodesmata and is prone to mislocalization artifacts from overexpression and fluorescent labeling. Hitherto, insight into PD ultrastructure has relied on electron microscopy (*19, 20*) and electron tomography (*21, 22*) of chemically fixed, heavy-metal-stained samples. These approaches revealed some features of PD architecture and formation, yet the required treatments denature biomolecules and compromise molecular interpretation (*23*). Recent advances in sample preparation and preservation have made cryo-electron tomography (cryo-ET) applicable to plant tissues (*24*). In cryo-ET, three-dimensional imaging of pristinely preserved samples at macromolecular resolution allows the visualization of membrane interfaces and reveals the organization of protein complexes in their native cellular context. In this study, we used cryo-ET to resolve the architecture of PD in the model plant *Physcomitrium patens* and describe mechanisms shaping the permeability of plant cell-cell interfaces.

### In situ architecture of plasmodesmata

While PD in land plants share a common evolutionary origin (*25*), their architecture varies across species and tissues. In vascular plants, like *Arabidopsis thaliana*, PD have diverse morphologies, including funnel-shaped and highly branched forms, reflecting tissue-specific adaptations (*26*–*29*). PD in non-vascular plants like the moss *Physcomitrium patens* remain less studied (*20, 30*). To investigate their PD architecture, we recorded tomograms of cell-cell junctions in the two dominant tissues of *P. patens*, protonemata and gametophores (Fig. 1). Unlike the morphological diversity observed in vascular plants, we exclusively found unbranched PD in *P. patens*, exhibiting canonical features: Each channel consists of a cell wall cavity lined by the plasma membrane, and traversed by a central tubular extension of the endoplasmic reticulum (ER), with a cytosol-filled intermembrane space, the cytoplasmic sleeve, between ER and plasma membrane (Fig. 1, movie S1 and S2). In wild type, PD length correlates with cell wall thickness, with thicker cell walls in gametophores necessitating longer PD (fig. S1). The neck regions, directly adjacent to the apertures at each end of the plasmodesmal channel, consistently represent the narrowest segments, imposing structural constraints on molecular passage. The tubulated ER within PD, also called the desmotubule, varies in diameter. In protonemata, desmotubules appear wider at neck regions (24.1 ± 2.2 nm) and narrower at the center of PD (10.5 ± 1.5 nm), while they remain uniformly wide (24.5 ± 3.1 nm) in gametophores. Desmotubules connect seamlessly to the cortical ER of adjacent cells, regardless of cortical ER morphology, which can appear as tubules (67%, top cell in Fig. 1, A and B, n = 61) with diameters substantially wider than the desmotubule inside PD or as sheets (33%, bottom cell in Fig. 1, C and D, n = 30) extending in parallel to the cell wall and cellular plasma membrane. Within the cytoplasmic sleeve, additional electron-dense material was observed. In protonemata, these likely proteinaceous features are most prominent at the neck regions, coinciding with wider desmotubule segments, but can also be detected along central desmotubule segments, where they are distributed sparsely. In gametophores, where desmotubules remain uniformly wide, these features are densely distributed along the entire length of PD. At narrow neck regions in both tissues, electron-dense material is most concentrated, obscuring desmotubules and the surrounding cytoplasmic sleeves.

**Fig. 1.**
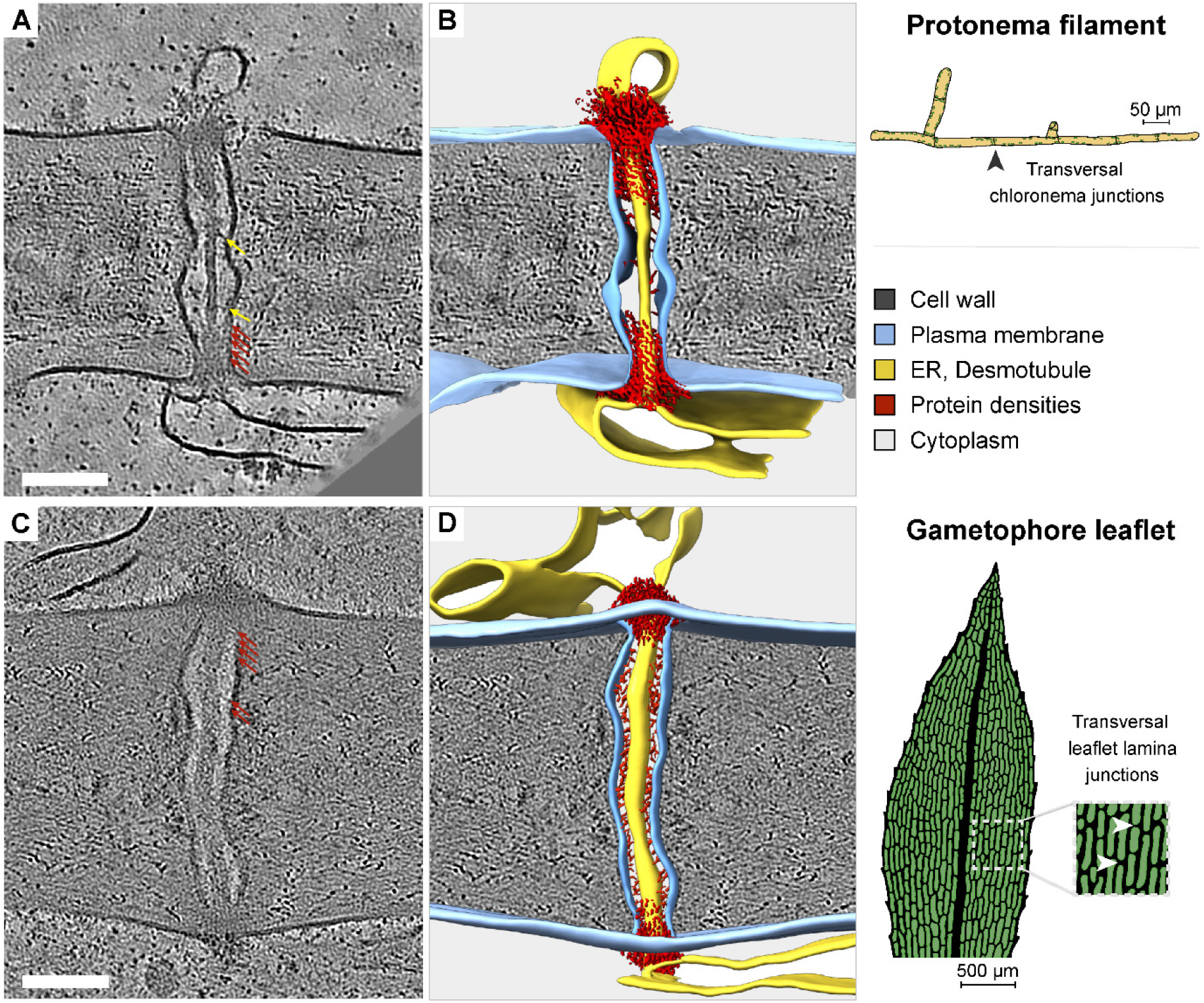
In situ architecture of plasmodesmata in *P. patens* revealed by cryo-ET. (**A**) 2D slices of a cryo-electron tomogram; cross-section of a cell wall between two cells within a protonema filament connected by a plasmodesma. Tethers connect wider desmotubule segments (red arrows) or narrower, central segments (yellow arrows) to the plasma membrane. (**B**) 3D-rendered segmentation of tomographic volume in A. (**C**) Slice through a plasmodesma connecting two gametophore leaflet lamina cells across their transversal cell wall interface. (**D**) Segmentation of the tomographic volume in C. Scale bars: 100 nm.

### Permeability control via callose deposition and degradation

To balance intercellular exchange and cellular homeostasis, plants dynamically regulate PD permeability. A key mechanism involves callose, a β-1,3-glucan cell wall polymer, which is deposited around neck regions by callose synthases and degraded by β-1,3-glucanases (*17, 31*–*33*). Synthesis and degradation either narrow or widen PD apertures, modulating permeability in response to developmental and environmental cues. Under conditions such as water scarcity, pathogen infection or during winter dormancy, plants induce callose synthesis via the stress-response hormone abscisic acid (ABA) (*34*–*37*). Unlike cellulose, which forms a dense fibrillar meshwork in the bulk cell wall, callose-rich deposits are structurally more amorphous (*38*). Cryo-ET of ABA-treated protonemata revealed large, granulated deposits of cell wall material around PD neck regions (Fig. 2, A and B). While neck regions generally became constricted after ABA treatment, 45% of PD (15 of n = 33) exhibited a more severe phenotype, where deposition led to complete abscission of cellular plasma membranes, ER, and cytoplasmic sleeves (movie S3). The cortical ER remained tethered to the plasma membrane near the former connection site to the desmotubule. These drastic changes in PD architecture provide a structural basis for the ABA-induced reduction in intercellular trafficking reported in *P. patens* (*39*). In contrast, we previously found that overexpression of the callose-degrading β-1,3-glucanase *Pp*GHL17_1 increases permeability of cell-cell interfaces (*11*). To determine whether and how increased permeability correlates with changes in PD architecture, we performed cryo-ET on a glucanase-overexpressing line, hereafter referred to as GHL17 (Fig. 2, C and D, and movie S4). We found PD in protonemata of GHL17 to be shorter and wider, not only around neck regions but along the entire channel (Fig. 2, E to H). Cell wall material with granulated texture at PD necks, weakly pronounced in WT and expanded upon ABA-treatment, is mostly absent in GHL17, supporting the interpretation that granulated textures correspond to callose-rich deposits. In accordance with PD conductance models predicting lower resistance and thus increased molecular flow in wider and shorter PD (*28, 40*), these architectural changes explain the enhanced cell-cell permeability phenotype in GHL17.

**Fig. 2.**
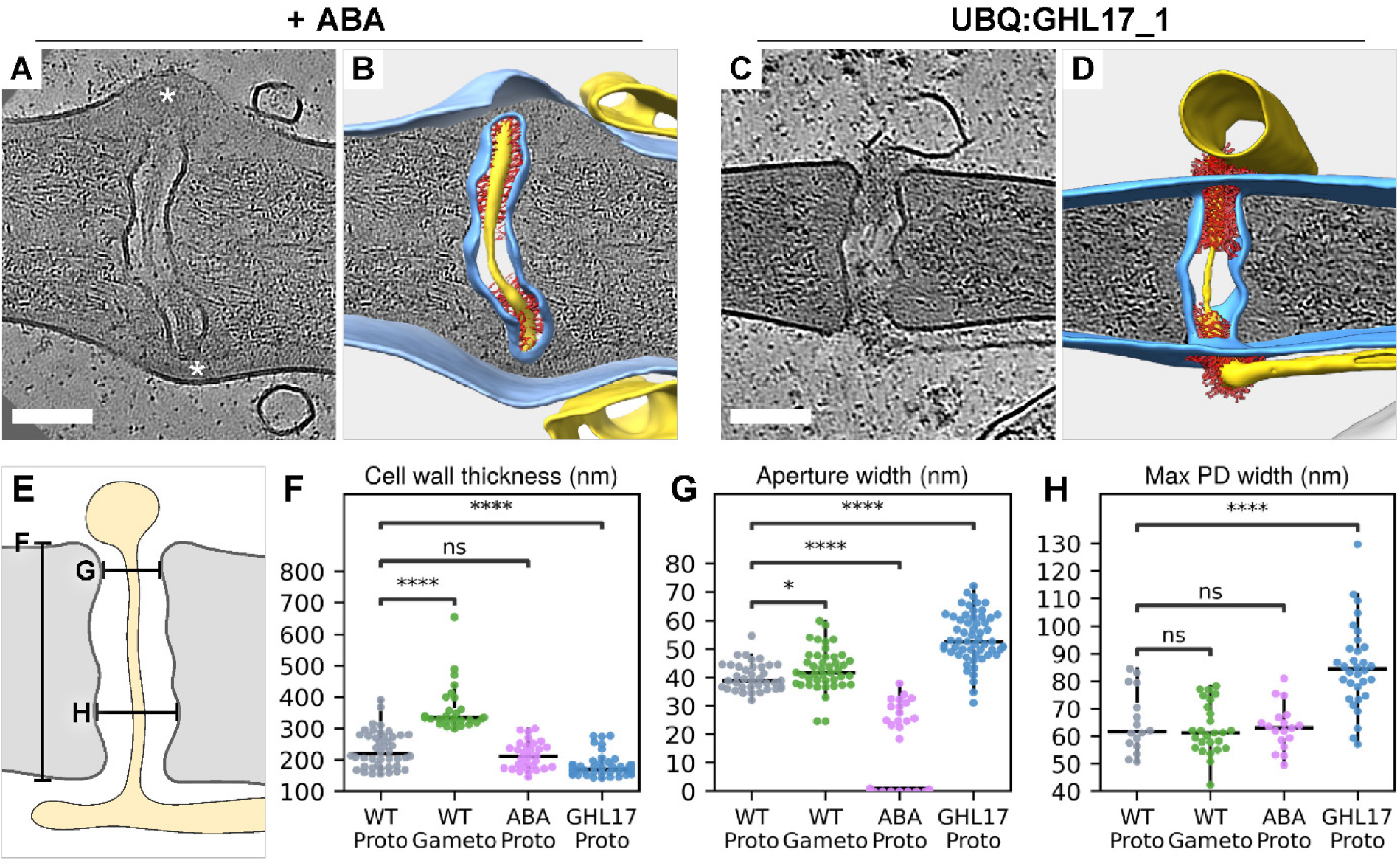
Deposition and degradation of the cell wall polymer callose gates plasmodesmata. (**A**) Incubating protonemata in 50 µM abscisic acid (ABA) for 17 h alters PD morphology. Asterisks mark putative callose deposits, disconnecting PD from cellular membranes and cytoplasm. (**B**) Segmentation of tomographic volume in A. (**C**) Overexpression of the callose-degrading enzyme GHL17 results in shorter and wider PD. (**D**) Segmentation of tomographic volume in C. (**E-H**) Effects of callose levels on PD architecture. (**E**) Schematic indicating measurement positions. (**F**) Quantification of cell wall thickness. (**G**) Aperture diameter at neck regions, representing the narrowest positions within PD. (**H**) PD diameter at the widest position. Mann-Whitney-Wilcoxon test (^*^P = 0.0252; ^****^P ≤ 0.0001; ns, not significant). Scale bars: 100 nm.

### Helical protein assemblies coat desmotubules at PD neck regions

Benefiting from wider PD in GHL17 protonemata, we identified periodic densities decorating all desmotubules at neck regions, while central segments lacked such features (Fig. 3A). These coatings cover approximately the first third of the desmotubule on both sides of PD, matching regions that appeared as wider desmotubules in WT protonemata. In GHL17, where cytoplasmic sleeves are widened, the electron-dense material that had obscured desmotubules at neck regions of WT PD now appeared to extend from the desmotubule coatings towards the plasma membrane. Prompted by the periodic appearance of the coatings, we performed subtomogram averaging along the desmotubule (fig. S2). At neck regions, we resolved assemblies of structurally repetitive units winding around the desmotubule and anchored to the membrane, forming a helical lattice composed of four intertwined wraps (Fig. 3B). The electron-dense material extending from the surface of the assemblies towards the plasma membrane escaped structure determination by averaging, consistent with a flexible arrangement. Similar helical assemblies were recovered by averaging desmotubules in WT protonemata (Fig. 3C), ABA-treated protonemata (Fig. 3D), and gametophores (Fig. 3E), indicating that they are defining architectural features of PD in *P. patens*, consistently present across different cell types and physiological conditions. Averaging of central desmotubule segments in GHL17 confirmed that these ordered coatings are restricted to PD neck regions in protonemata (Fig. 3F). Assembly lengths varied across tissues and conditions. In gametophores, the entire desmotubule is coated, whereas in protonemata, assemblies are confined to the neck region but vary in length (Fig. 3I). Their repetitive architecture and variable lengths suggest a modular organization, assembled from recurring structural units.

**Fig. 3.**
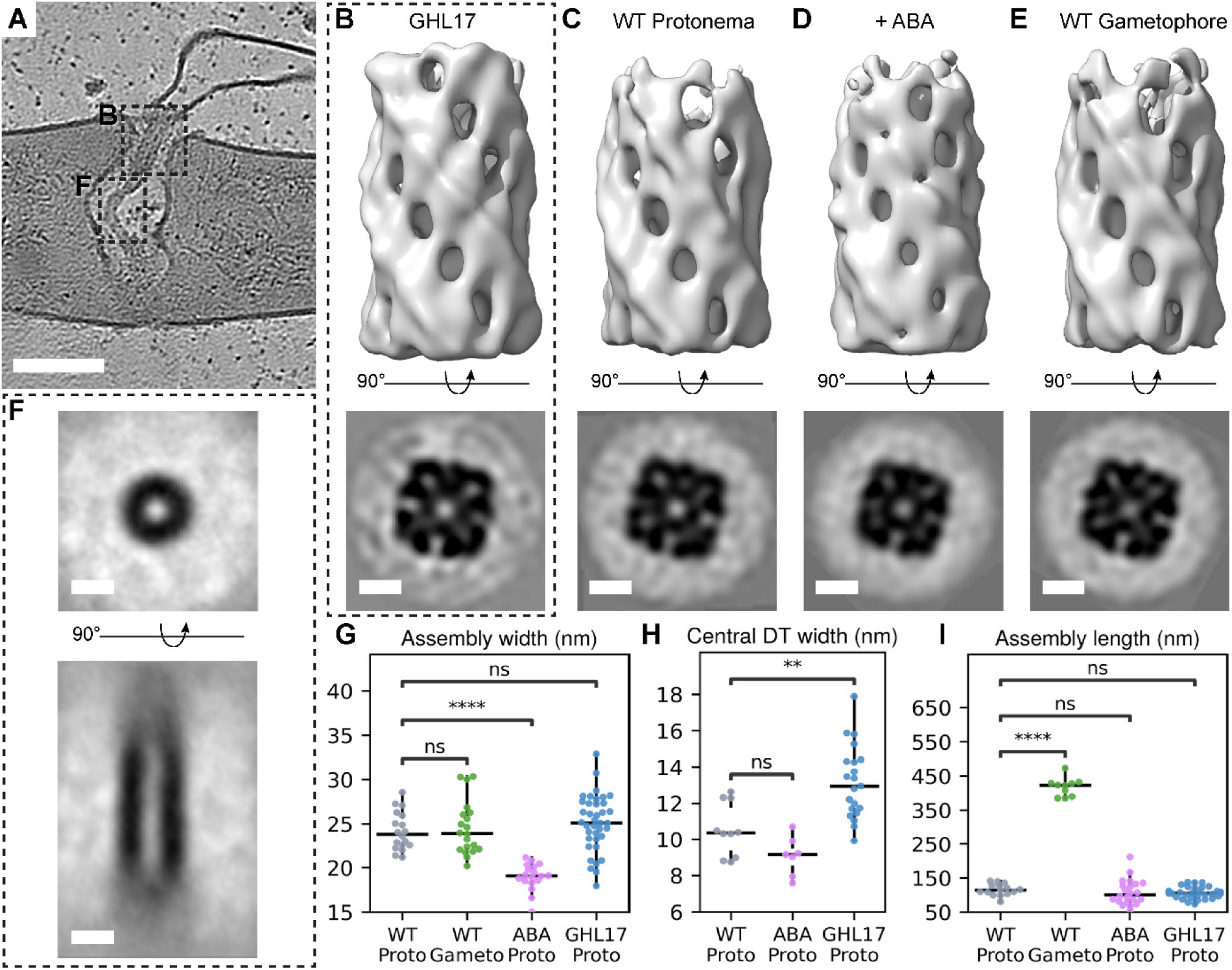
Helical desmotubule-coating assemblies are core architectural features of plasmodesmata. (**A**) Tomographic slice through a plasmodesma in GHL17 protonema. Dotted rectangles exemplify regions where subtomograms for averaging along the desmotubule were extracted. (**B-E**) Resolved desmotubule-coating assemblies in (**B**) GHL17 protonemata, (**C**) WT protonemata, (**D**) ABA-treated WT protonemata and (**E**) WT gametophores. Top panel: Side view of 3D-rendered assemblies. Bottom panel: 2D cross-sections through assemblies. (**F**) Transverse (top) and longitudinal (bottom) cross-sections of the averaged central desmotubule segment in protonemata, where assemblies are absent. (**G-I**). In situ measures of assemblies. (**G**) Assembly width. (**H**) Diameter of central desmotubule segment in protonemata. (**I**) Assembly length. Mann-Whitney-Wilcoxon test (^**^P = 0.0019; ^****^P ≤ 0.0001; ns, not significant). Scale bars: (A) 100 nm; (B-F) 10 nm.

### MCTPs coat the desmotubule and tether it to the plasma membrane

To elucidate the molecular composition of desmotubule-coating assemblies, we derived a list of candidate proteins from a proteome of PD-enriched cell wall fractions ((*11*), and table S1). Proteins predicted to localize to the ER and those with potential scaffolding functions were included, while excluding those assigned to the secretory pathway. We predicted structures of the remaining 43 candidate proteins and generated synthetic density maps for comparison with experimental maps of the assembly. Among all tested candidates, exclusively Multiple C2 domain and Transmembrane Proteins (MCTPs) matched the experimental data (Fig. 4, fig. S3, and movie S5). MCTPs co-localize with PD markers in land plants and have been identified in proteomes of PD fractions across species (*6*–*13*). In *P. patens*, the MCTP family includes six homologs (*Pp*MCTPs, table S2 and fig. S4) with conserved domain topology, three of which are highly enriched in PD fractions (*Pp*MCTP1, 2, and 5). MCTPs are ER-associated membrane proteins containing a C-terminal reticulon homology domain (RHD). RHDs are widely conserved across eukaryotes and function in ER shaping (*41, 42*), including in ER-containing cytoplasmic bridges in the insect germline (*43*). Through structure prediction of multiple oligomeric configurations, we identified dimers as the most likely arrangement of *Pp*MCTPs, with interfaces formed by two central α-helices per monomer and their membrane-embedded RHDs (fig. S4 and fig. S5). Given the high sequence conservation among *Pp*MCTPs (77% global identity, 94% at the dimer interface), heterodimerization appears likely. Predicted homo-and heterodimers are nearly identical in overall structure, differing primarily in the length of a glycine-rich linker connecting the N-terminal C2A domain to C2B (C2A-B linker domain), which ranges from 80 to 150 residues across family members (fig. S4 and table S2). In total, *Pp*MCTPs contain four C2 domains. C2B, C2C, and C2D are positioned around the central dimer interface, while C2A is flexibly tethered via the extended C2A-B linker domain (Fig. 4A). To determine how MCTPs organize within the desmotubule-coating assembly, we excluded the positionally flexible C2A and its linker and fitted the truncated dimers into the density map obtained by subtomogram averaging. These truncated dimer units fully account for the resolved helical assembly coating desmotubules in situ (Fig. 4B and fig. S3), affirming MCTPs as its fundamental building blocks. Although side chain interactions remain unresolved at this resolution, we identified potential contact sites of individual dimer units at the domain-level. In the resolved assembly, C2B domains interlace with C2B domains of adjacent dimers in the neighboring helical wrap (Fig. 4C), while C2C and C2D domains contact adjacent dimers within the same wrap (Fig. 4D). The ability of C2 domains to mediate protein-protein interactions has been reported for other C2-containing proteins (*44, 45*). Deletion of C2C in *Arabidopsis thaliana* MCTP4 (*At*MCTP4) disrupts PD association (*22*), supporting its role in stabilizing MCTP assemblies at PD. MCTPs have recently been described to prevent ER membrane abscission at the developing cell plate during cytokinesis, thereby facilitating PD formation through incomplete cell division (*22*). The assemblies resolved here may provide the structural basis for this function, stabilizing the desmotubule and shielding it against membrane severing (Fig. 4E). Supporting this, we observed that under ABA-induced callose deposition, MCTP assemblies remain intact and are internalized into the cell wall along with desmotubules, plasma membrane tethers and cytoplasmic sleeves (Fig. 2, A and B). Severing occurs cytoplasm-proximal to the assembly, consistent with a protective role against membrane abscission during both cytokinesis and in mature PD.

**Fig. 4.**
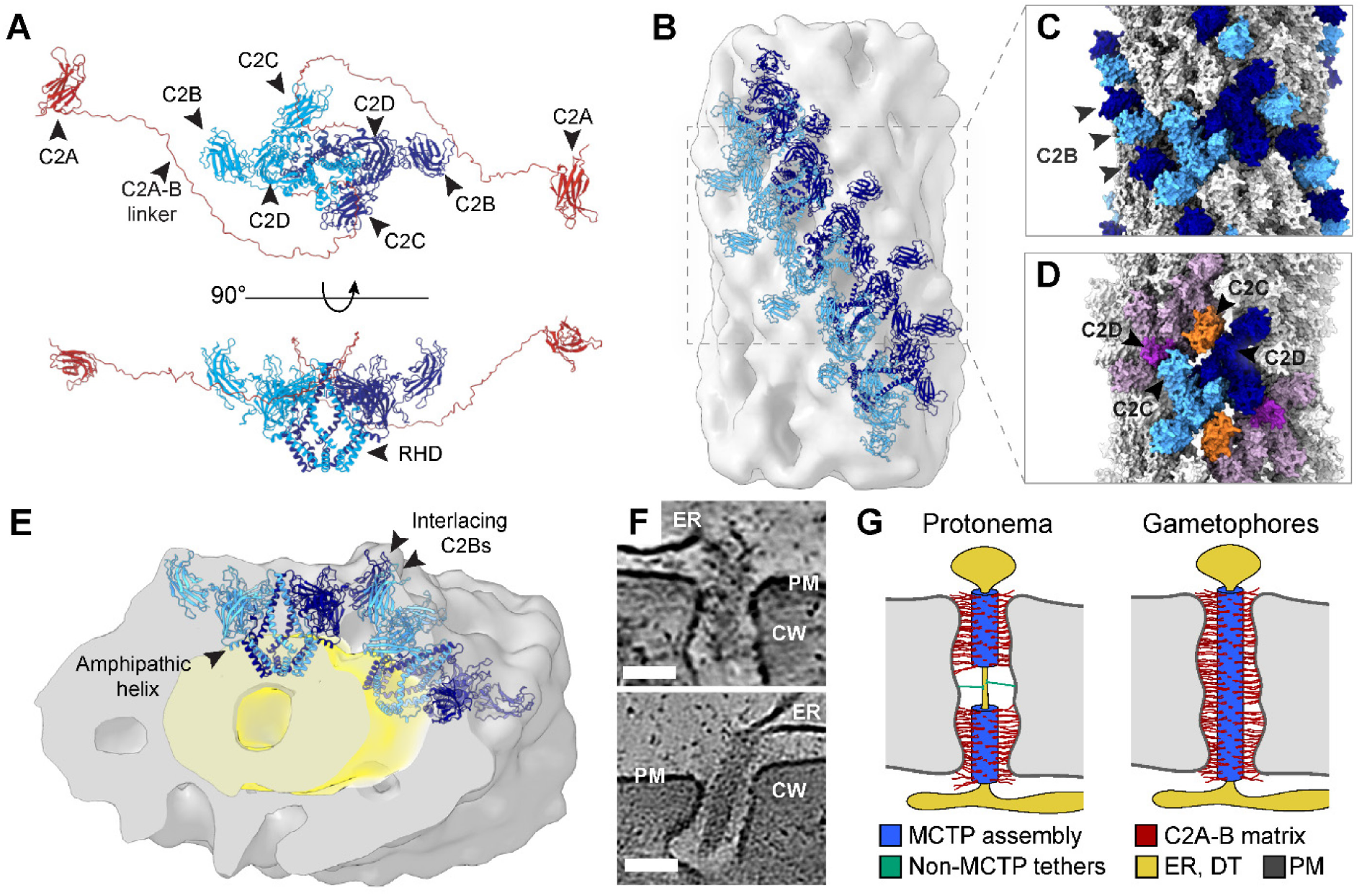
Arrangement of *Pp*MCTPs in the desmotubule-coating assembly. **(A)**.Structure prediction of full-length, dimeric *Pp*MCTP. N-terminal C2A domains connect through flexible linkers (red) to the dimerizing core (monomers in light and dark blue). Four central α-helices form the dimer interface. Reticulon homology domains (RHDs) anchor the dimer to the ER-derived desmotubule membrane. C2B, C2C and C2D domains are accessible for protein-protein interactions. **(B)**.Subtomogram average of the desmotubule-coating assembly with docked models of dimeric *Pp*MCTP (blue), tiled along one of four helical wraps. Flexible linkers and C2As are not resolved in average. (**C**) Surface rendering of the assembly. C2B domains of dimers from adjacent helical wraps interlace (light and dark blue). (**D**) Dimers within the same helical wrap contact via C2C and C2D domains. (**E**) Angled cross-section of the assembly (grey) with two docked dimers (blue) and desmotubule membrane region (yellow). (**F**) Magnified views of *Pp*MCTP assemblies in situ. Scale bars: 50 nm. (**G**) Schematic of desmotubule organization in protonemata and gametophore tissues. MCTP assemblies (blue) coat desmotubules (DT), with C2A domains and C2A-B linkers (red) extending towards the plasma membrane (PM). In protonemata, central desmotubule regions are not coated by MCTP assemblies, with occasional tethers extending to the plasma membrane (green). In gametophores, desmotubules are fully coated by MCTP assemblies.

### Extended C2A domains enable desmotubule-plasma membrane tethering

In mature, post-cytokinetic cells, MCTPs contribute to permeability control through a callose-independent mechanism (*16*). Higher-order mutants of *At*MCTPs exhibit increased intercellular flux, and molecular dynamics simulations suggest that the C2B, C2C, and C2D domains of *At*MCTP4 can interact with PIP4 lipids in the plasma membrane. This lipid-mediated tethering has been proposed to modulate cytoplasmic sleeve diameter and thereby influence permeability independently of callose turnover (*16*). While we cannot exclude membrane binding of these domains, we did not observe direct contacts between the plasma membrane and the surface of the resolved fraction of *Pp*MCTP assemblies, containing C2B, C2C, and C2D domains. However, with theoretical lengths of 30 - 50 nm, fully extended C2A-B linkers are sufficiently long to bridge the cytoplasmic sleeve, allowing C2A binding at the plasma membrane. We therefore measured the distance between the assembly surface and the plasma membrane in GHL17 plasmodesmata and found a maximum tether span of 26.4 nm (19.8 ± 4.2 nm, n = 72, fig. S1F). While C2A domains and their linkers are too small and flexible to be resolved by subtomogram averaging, we observed electron-dense material in individual tomograms extending from the assembly surface toward the plasma membrane, coinciding with plausible trajectories of extended C2A-B linker domains (Fig. 4F). These densities terminate at membrane-proximal positions, consistent with C2A-mediated lipid binding, supporting a model in which C2A domains serve as the primary tether linking desmotubules to the plasma membrane in *P. patens*. In mature gametophores, where assemblies and tethers span the full PD length, reduced intercellular flux has been reported (*30*), while loss of MCTPs in Arabidopsis increases cell-to-cell conductivity (*16*). These findings illustrate that MCTP levels correlate with PD permeability. Models invoking phase-separated protein networks as molecular basis for selective permeability, analogous to FG-NUPs in the nuclear pore, have been discussed for PD (*13, 46, 47*). Upon closer inspection, C2A-B linker domains exhibit sequence features consistent with a role in permeability control via phase separation based on electrostatic rather than hydrophobic interactions. The C2A-B linker domains are polyampholytic - that is, they contain a balanced distribution of oppositely charged residues interspersed with flexibility-promoting amino acids (fig. S4), enabling transient electrostatic interactions with other charged macromolecules (*48*). C2A-B linker domains are further predicted to be highly disordered and to exhibit strong intrinsic propensity for spontaneous liquid-liquid phase separation (LLPS propensity > 93%, fig. S6). Despite high overall conservation across the MCTP family, C2A-B linker domains are the most variable sequence regions, hinting at potential functional diversification. Given these properties, we propose that C2A-B domains, likely in interaction with proteins extending from the plasma membrane, form a mesh-like, selective barrier within the cytoplasmic sleeve to modulate diffusion across PD. As MCTP assemblies are constitutively present, whereas sleeve diameter at neck regions is shaped by dynamic changes in callose levels, both mechanisms may act synergistically. Low callose levels widen the sleeve, stretching linkers and loosening the mesh, whereas high callose levels constrict the sleeve, concentrating proteins and tightening the barrier.

Altogether, our data indicate that MCTP dimers form helical assemblies coating the desmotubule, anchored via RHDs which contribute to membrane tubulation. Assemblies are stabilized by protein-protein interactions at C2B, C2C and C2D domains. C2A domains tether the desmotubule to the plasma membrane, while polyampholytic C2A-B linkers populate cytoplasmic sleeves, potentially forming a dynamic diffusion barrier (Fig. 4G).

## Conclusions

In this work, we report native architectures of plasmodesmata across tissues and physiological states in *P. patens*, providing molecular context for how cell wall, membrane and protein components are arranged to shape intercellular connectivity in plants. Callose deposition narrows the apertures, acting as an apoplasmic valve capable of fully sealing PD. MCTPs, previously implicated in primary PD formation and permeability modulation, form membrane-anchored helical assemblies that coat and stabilize the desmotubule while tethering it to the plasma membrane. MCTP assemblies are retained in mature PD, where they preserve desmotubule organization and maintain ER-plasma membrane contact sites, even when symplasmic membrane continuity is lost due to callose-induced closure. Our findings indicate MCTP assemblies as ubiquitous architectural constituents of PD and identify membrane abscission as a structural outcome of callose-dependent closure in post-cytokinetic interfaces.

## Acknowledgements

We thank D. Morado, T. Schäfer, D. Bollschweiler, C.O.J. Kaiser, O.H. Schiøtz, P.S. Erdmann, and the J. Briggs group for access, training and outstanding support with TEM and FIB instruments. We are grateful to A. Bieber and C. Capitano for their expert guidance in cryo-FLM methodology. F. Beck, A. Bieber, M. Riggi and M. Weberskirch provided invaluable input on data processing and visualization. We thank N.S. Kazemein Jasemi for providing the GHL17 cell line; the Max Planck Computing and Data Facility (MPCDF) for HPC support; and the B.D. Engel and M.M. Wudick groups for fruitful discussions and feedback along the way.

