## Supplemental Materials for "In situ architecture of plasmodesmata suggests mechanisms controlling intercellular exchange"

### Materials and Methods

Preparation of Physcomitrium patens tissues for cryo-ET

Culturing, sample preparation, cryo-fixation, and FIB milling procedures are described in detail in Pöge et al. (*24*). Briefly, *P. patens* (Hedw.) Gransden ecotype was grown at 25 °C on BCDAT agar medium overlaid with cellophane, under 18 h white light / 6 h dark cycles. Small protonemata patches were detached using tweezers, submerged in freezing buffer (BCDAT liquid medium supplemented with Ficoll 400 dissolved in $H_2O$ to a final concentration of 20% [w/v]), and immediately high-pressure frozen (LEICA EM ICE, Leica Mikrosysteme GmbH, Wetzlar, Germany). Gametophores were collected after 4 weeks by manually dissecting individual phyllids with fine scissors, followed by immediate immersion in freezing buffer and high-pressure freezing. Samples were vitrified using both the "waffle method" and freezing in 3 mm carriers with a 100 µm deep cavity, optimized for subsequent Serial Lift-Out FIB milling.

Cryo-ET: Data acquisition

Tilt series were collected using a Titan Krios G3i instrument at 300 kV, equipped with a Selectris X energy filter and a Falcon 4i camera (Thermo Fisher Scientific). Tilt series were recorded using the Tomography5 software package (Thermo Fisher Scientific). A dose-symmetric tilt scheme was used with an angular increment of 3°. Magnification was set to 64,000x (pixel size = 1.89 Å) and a dose rate of 3 $e^-/Å^2$ per tilt, resulting in a total dose of 100 $e^-/Å^2$ per tilt series. The target defocus ranged from -3.0 to -6.0 µm. A small dataset was collected on a Titan Krios G2 equipped with a Gatan post column energy filter and K2 Summit (Gatan) camera. Here, tilt series were recorded at a magnification of 42,000x (pixel size = 3.52 Å) using the SerialEM software (*49*).

Tomogram reconstruction and segmentation

Tilt series were processed as described in Pöge et al. (*24*). Preprocessing including motion correction, removal of bad tilt images and dose filtering were performed in TOMOMAN (version 0.6.9, (*50*)). If tilt-series' contained at least 5 platinum fiducials evenly distributed over the field-of-view, IMOD (version 4.11.25, (*51*)) fiducial tracking was employed for tilt-series alignment. Otherwise, AreTomo (version 1.3.3, (*52*)) was used for tilt-series alignment. Afterwards, 4x binned tomograms were reconstructed with the weighted back projection algorithm implemented in IMOD. Membranes were segmented with MemBrain-Seg (*53*). While this software reliably traces intracellular membranes, it barely picked plasmodesmata membranes. Hence, the latest built-in model (v10) was refined following the instructions available at https://teamtomo.org/membrain-seg/Usage/Training/. Furthermore, tilt-series were imported into WARP (version 1.0.9, (*54*)). Membrane tethering densities within cytoplasmic sleeves were segmented using a convolutional neural network (CNN) trained in EMAN 2.99 (*55*). Because these features span only 1-2 pixels in width, training and segmentation were performed on bin2 tomograms which were denoised with cryo-CARE (*56*) and subsequently processed with IsoNet (*57*) employing the default parameters. For each tomogram, 1000-1200 negative reference regions were selected from areas containing cell wall, cytosol, or non-PD membranes. Additionally, 30-50 positive reference areas were selected per tomogram, representing regions along the full PD channel. Positive references were manually annotated, serving as training input (training parameters: learnrate = 0.0001; iterations = 200; ncopy = 5;

batch = 20; nkernel = 40,40,1; ksize = 9,7,5; poolsz = 2,1,1, box size = 128). While picking up tethers within cytoplasmic sleeves, the network failed to identify individual tethers at highly dense PD aperture regions. For those regions an additional voxel density thresholding step was performed in Amira (Thermo Fisher Scientific). All tomograms displayed in this work were denoised with cryo-CARE and visualized with IMOD (version 4.11.25). Segmentations were curated in Amira and jointly rendered in ChimeraX (*58*).

Cryo-ET: Tomogram analysis

PD dimensions were manually annotated with the contour tool in 3dmod and the corresponding distances calculated in MATLAB (version 2022a). Data visualization and statistical analysis were performed in Python using the libraries matplotlib, numpy, csv, pandas, seaborn, scipy, and statannotations. Statistical comparisons were conducted using two-sided Mann-Whitney-Wilcoxon tests, implemented via the statannotations.Annotator class.

Cryo-ET: Subtomogram averaging of the desmotubule-coating assembly

For each desmotubule-coating assembly, a two-point contour was manually defined in 3dmod. The first point marked the beginning of the complex within the PD channel, and the second was placed at the channel opening toward the cytosol. In gametophore samples containing a single continuous assembly spanning the entire PD, a single long contour was defined from one channel opening to the other. Subtomograms (4× binned, voxel size = 7.6 Å, box size = 64 voxels) were extracted at 1 nm intervals along the line connecting these two points. Euler angles for θ (Theta) and ψ (Psi) were calculated such that the z-axis of each subtomogram aligned parallel to the desmotubule contour axis; φ (Phi) angles were randomized. Initial alignment was performed in STOPGAP (v0.7.0, (*59*)) with shifts only allowed perpendicular to the axis of the PD channel without φ angle search. The reference for the first iteration was the average of the extracted subtomograms without alignment (fig. S2). Subsequently, the average of the current iteration was used for the next one. All references were filtered to a resolution of 35-40 Å.

For the GHL17 dataset, pseudo-symmetry expansion and a second alignment step in STOPGAP were performed. From every initial subtomogram position after the first alignment step, 11 new positions were defined by shifting 6 nm radially outward onto the desmotubule surface with an angular distance of ~33°. Euler angles were updated to align the subtomogram z-axis perpendicular to the desmotubule surface using the STOPGAP script sg_motl_shift_and_rotate.m. Subtomograms were re-extracted at these oversampled positions (4× binned, voxel size = 7.6 Å, box size = 48 voxels) and aligned with exhaustive search over all three Euler angles, allowing a maximum shift of 11 nm. Following alignment, subtomograms were distance-filtered by removing particles within 8 nm of one another, retaining only the subtomograms with the highest alignment score (fast local correlation function in STOPGAP). Subsequently, subtomograms were extracted in WARP (2× binned, voxel size = 3.8 Å, box size = 96 voxels) and subjected to 3D classification, refinement, and postprocessing in RELION (v3.0.5, (*60*)). Resolution of the GHL17-derived average was estimated according to the Fourier Shell Correlation ($FSC_{0.143}$ ≈ 33 Å, fig. S2).

To enable comparison of desmotubule-coating assembly across all datasets, subtomograms were extracted in WARP (voxel size = 7.6 Å, box size = 64 voxels) directly after the initial alignment in STOPGAP without symmetry expansion. These subtomograms were classified and aligned in RELION. During postprocessing, all averages were filtered to a resolution of 40 Å, enabling comparison (Fig. 3, B to E).

Proteome-derived list of desmotubule-coating candidates

The *Physcomitrium patens* high-confidence PD proteome (HC300; (*11*)) was filtered for plasma membrane and endoplasmic reticulum-localized proteins containing at least one transmembrane or known membrane-binding domain. In addition, proteins annotated with potential scaffolding functions, as well as those lacking functional annotation in *P. patens*, were included. Transporters, hydrolytic enzymes, proteins with known functions in vesicle transport, and proteins predicted to be secretory pathway-targeted were excluded. This filtering yielded 43 candidate proteins for further assessment (Supplementary Table S1).

AlphaFold predictions of candidate proteins

Monomeric structures of all candidate proteins were predicted using AlphaFold v2.3.1 (*61*). While monomers could in principle fit within the experimental density, their small size allowed a wide range of orientations, often resulting in biologically implausible placements (e.g., membrane-associated domains facing the cytoplasmic sleeve) or complete burial within the map without accounting for key structural features. These ambiguities precluded reliable identification of monomeric fits. To address this, each protein was further evaluated individually using structural, biochemical, and localization information from the literature, with the goal of identifying higher-order oligomers from a combination of monomeric PD proteins that could recapitulate the desmotubule-coating assembly. For proteins with documented oligomeric states, dimeric and trimeric structures were predicted using AlphaFold Multimer v2.3.1 (*62*). Predicted complexes with high confidence (0.8*ipTM + 0.2*pTM ≥ 0.60) were subjected to rigid-body fitting into the experimental density. For models containing extended disordered regions (pLDDT < 50), truncated variants retaining only structured domains and short interdomain linkers were generated. Non-MCTP candidates failed to align meaningfully with the density, producing fits that either occupied the volume nonspecifically or conflicted with membrane topology. Adding repeating units of non-MCTP oligomers did not recover the observed organization of the coat (fig. S3A). In contrast, truncated *Pp*MCTP dimers aligned closely with the experimental density, recapitulating the periodic features of the desmotubule-coating assembly (fig. S3, B and C). To quantify the fit, we generated a full model by tiling *Pp*MCTP dimers using rigid-body fitting, simulated a corresponding density map from the atomic model at the resolution of the experimental average, then calculated the cross-correlation between the synthetic and experimental map (fig. S3D).

Assessment of MCTP structure by structure predictions

To explore the structural basis of MCTP oligomerization, we predicted monomeric, dimeric and trimeric structures for each of the six *Pp*MCTPs using AlphaFold v2.3.1 and AlphaFold Multimer v2.3.1. Dimeric assemblies were further modeled for all pairwise combinations of *Pp*MCTPs to evaluate the plausibility of homo- vs. heterodimerization (fig. S4C). Only predictions with a weighted score of *0.8*ipTM + 0.2*pTM ≥ 0.6* were considered. Resulting structure predictions were further assessed based on pLDDT and PAE scores. For trimer predictions (fig. S4B and S5), the flexible N-terminus was truncated to reduce compute. AlphaFold3 prediction was performed on https://alphafoldserver.com/ (*63*).

In silico profiling of sequence-encoded biophysical properties of full-length MCTPs

We first noticed C2A-B linker regions for their high glycine content at the sequence level. In subsequent structure predictions, they consistently scored below 50 in pLDDT, by now a well-established indicator of intrinsic disorder (*64*). To further characterize biophysical C2A-B properties, we applied prediction algorithms specifically aimed to assess disorder and phase separation promoting properties (fig. S6). FuzDrop estimates propensity of amino acid sequences for droplet-formation (pDP) and liquid-liquid phase separation potential (pLLPS) (*65*). We also applied AIUPred, a neural network-based successor of IUPred, which improves disorder prediction accuracy through deep learning (*66*). pLDDT scores from AlphaFold2-Multimer (ranging from 1-100) were normalized by dividing by 100 to match the 0-1 scale of pDP and AIUPred.

### Supplementary Information

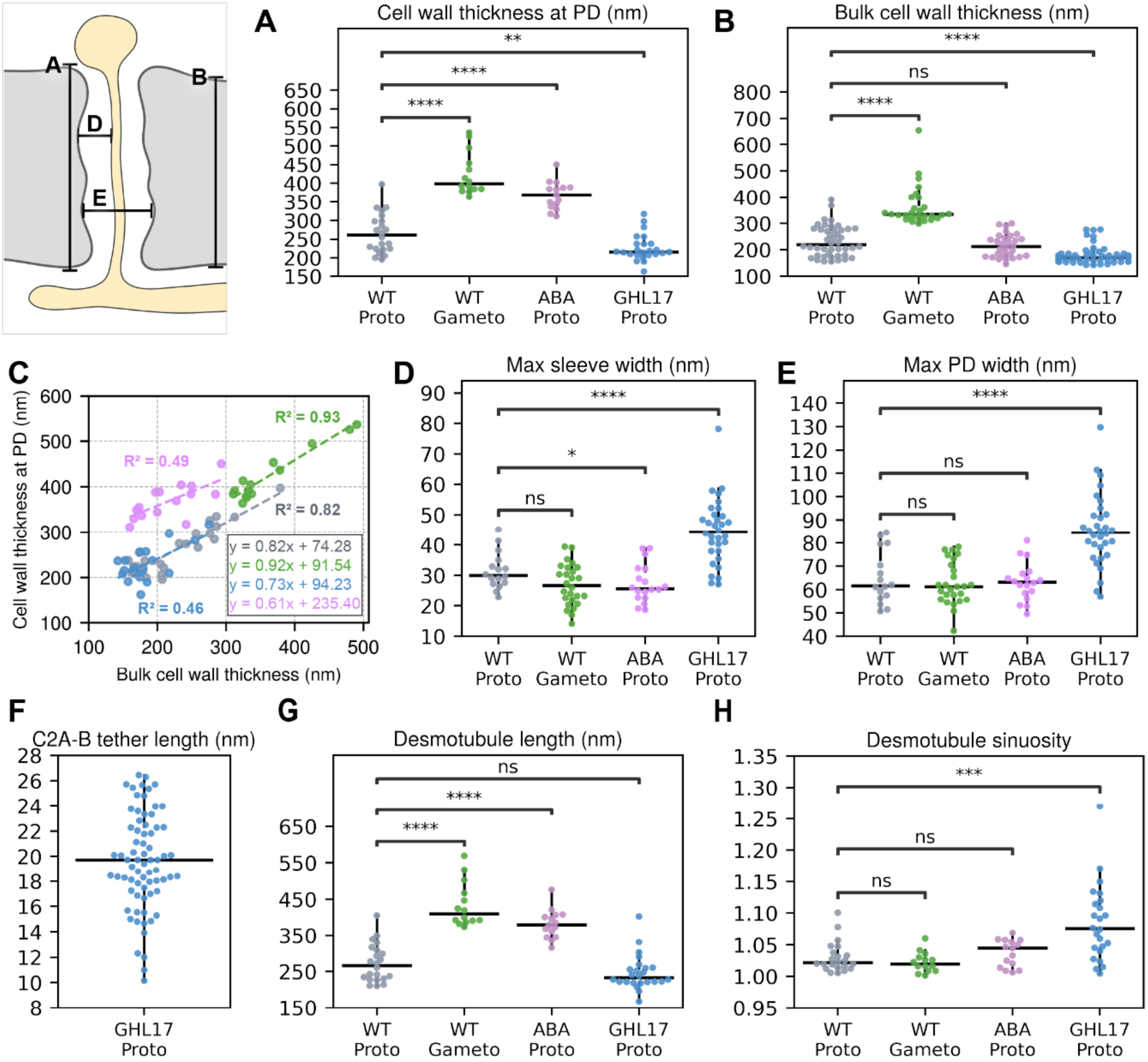


**Fig. S1. In situ measurements.**
Quantification of plasmodesmata features in wild type protonemata (WT Proto, grey), wild type gametophores (WT Gameto, green), ABA-treated wild type protonemata (ABA Proto, magenta), and UBQ:GHL17_1 protonemata (GHL17 Proto, blue). **(A)** Cell wall thickness at plasmodesmata. For sealed plasmodesmata in ABA-treated tissue, end-to-end length including callose plugs was measured. **(B)** Bulk cell wall thickness measured in regions away from plasmodesmal channels, excluding bulged regions at plasmodesmata necks. Short-term ABA treatment (50 µM for 17h) specifically induces bulging at plasmodesmata neck regions, whereas constitutive GHL17 overexpression broadly affects cell wall thickness at and away from plasmodesmata. **(C)** Scatter plot of cell wall thickness at PD (approximating PD length) versus bulk cell wall thickness with linear regression fits per data set (dashed lines); slopes and $R^2$ (coefficient of determination) values indicate the degree and consistency of coupling. Lower $R^2$ in ABA and GHL17 data reflects increased local cell wall heterogeneity compared to WT. **(D)** Maximum width of cytoplasmic sleeve between desmotubule and plasma membrane. **(E)** Maximum width of plasmodesmal channels, measured from plasma membrane to plasma membrane. **(F)** Length of tethers originating from surfaces of desmotubule-coating assemblies and terminating at the plasma membrane. **(G)** Total length of desmotubules measured along their curvature. **(H)** Sinuosity of desmotubules as an indicator of curvature. Desmotubules in GHL17 plasmodesmata have lengths similar to those in wild type but are accommodated within shorter channels, resulting in increased sinuosity. Mann-Whitney-Wilcoxon test (*P = 0.04; **P = 0.008; ***P = 0.0004; ****P ≤ 0.0001; ns, not significant).

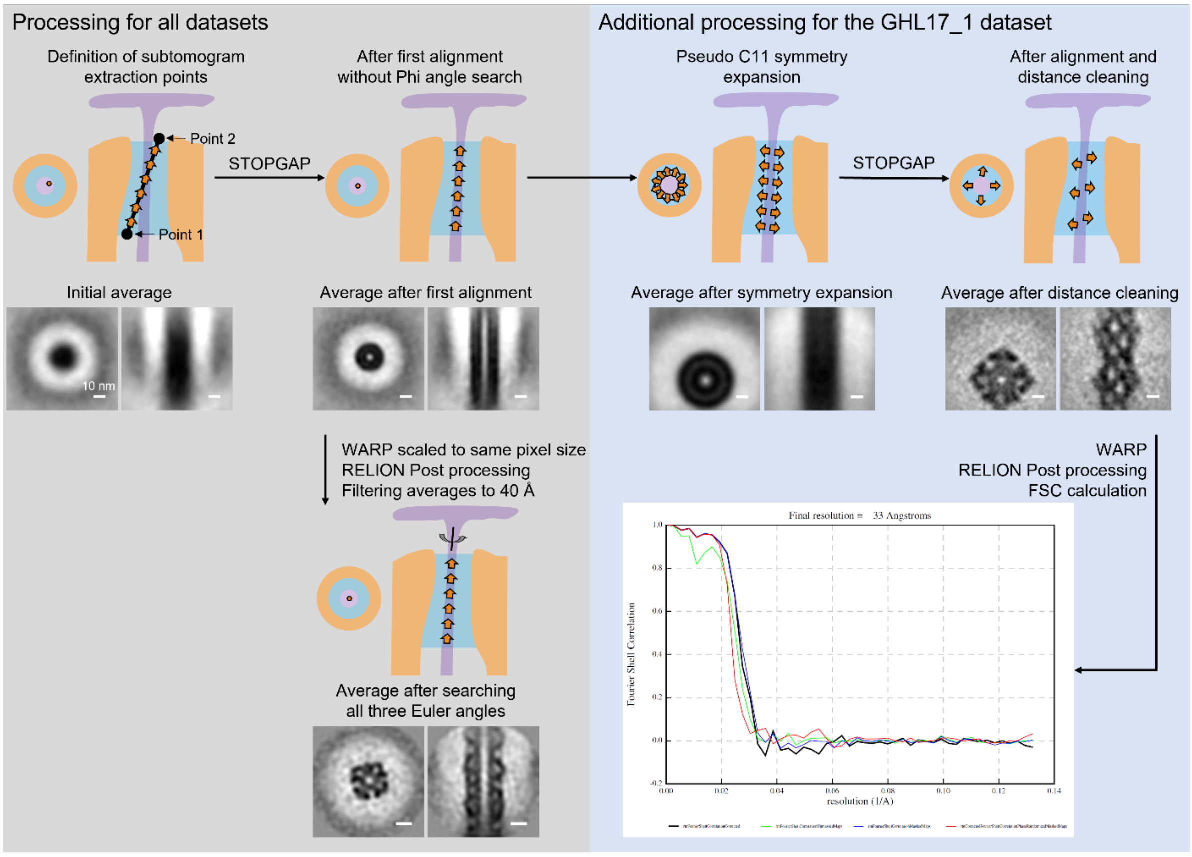


**Fig. S2. Subtomogram averaging workflow.**
For each alignment step, the top panel shows a schematic representation, and the bottom panel displays an xy cross-section and an xz-plane side view through the center of the resulting subtomogram average. Subtomogram extraction points are indicated by orange arrowheads. The direction of each arrow represents the subtomogram z-axis as defined by the Euler angles θ (Theta) and ψ (Psi). Scale bars: 10 nm.

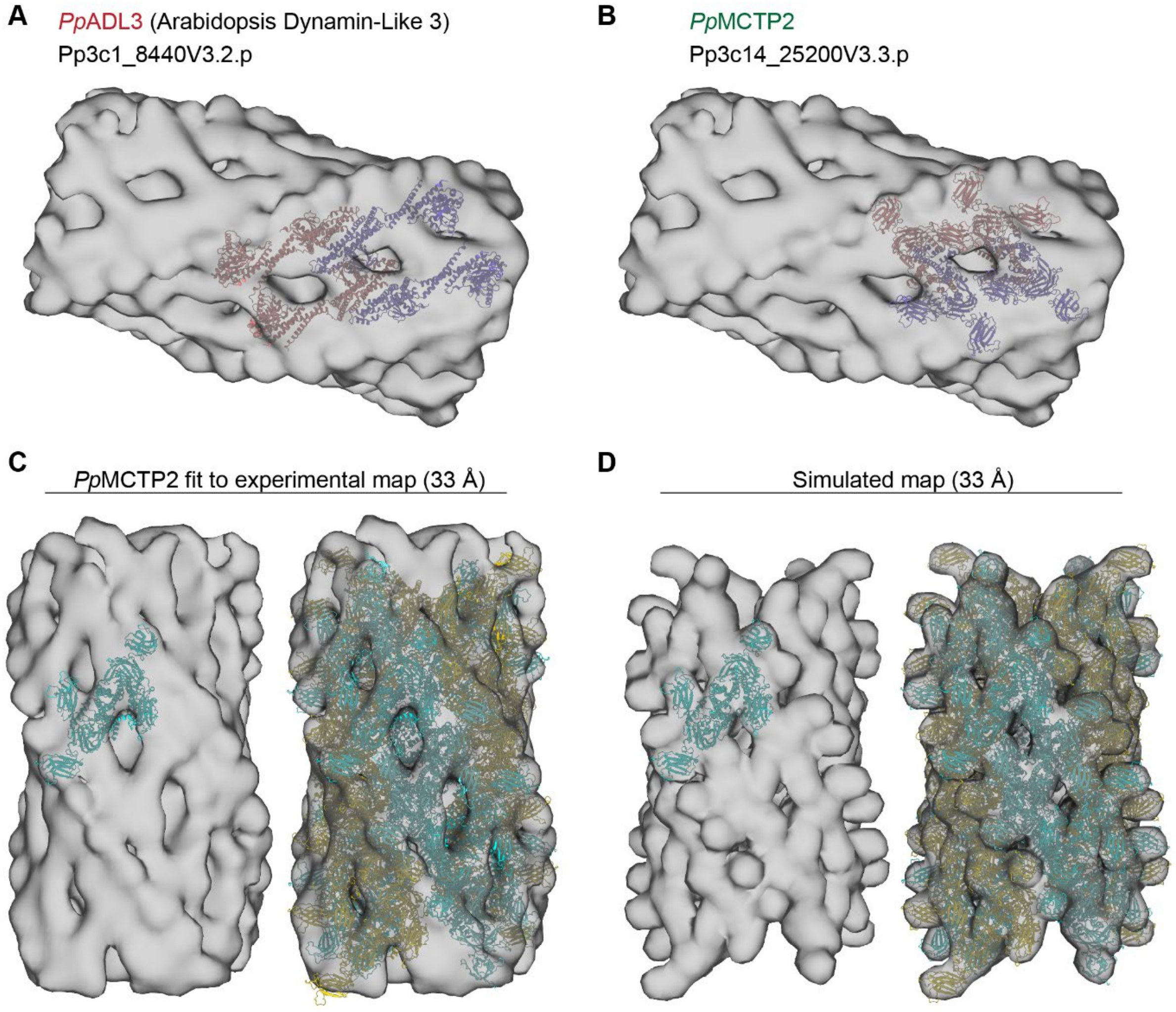


**Fig. S3. Fitting of predicted candidate protein structures into experimental density map.**
To evaluate potential components of the desmotubule-associated coat, we tested candidates identified in PD-enriched proteomes. Dynamin-related proteins emerged as plausible candidates based on their property to assemble into large complexes associated with ER and their membrane tubulating function (*67*). Such assemblies are formed by repeating dynamin dimers. (**A**) Two predicted *Pp*ADL3 dimers were fitted into the experimental density map of the desmotubule-coating assembly (monomers in pink and purple). Although the dimer models span the assembly lumen, the fit leaves substantial regions unaccounted for and fails to recapitulate key structural features. (**B**) In contrast, *Pp*MCTP2 dimers align closely with the experimental map, matching local geometries. (**C**) A full molecular model was generated by tiling *Pp*MCTP2 dimeric core units along the resolved coat. Dimers colored in cyan and yellow, alternating colors between helical wraps. (**D**) From the full molecular model based on the experimental map, we generated a corresponding synthetic map. Filtered to 33 Å resolution, the synthetic map closely resembled the experimental density map. The cross-correlation coefficient between synthetic and experimental map is 0.92, indicating a good overall agreement between model and experiment.

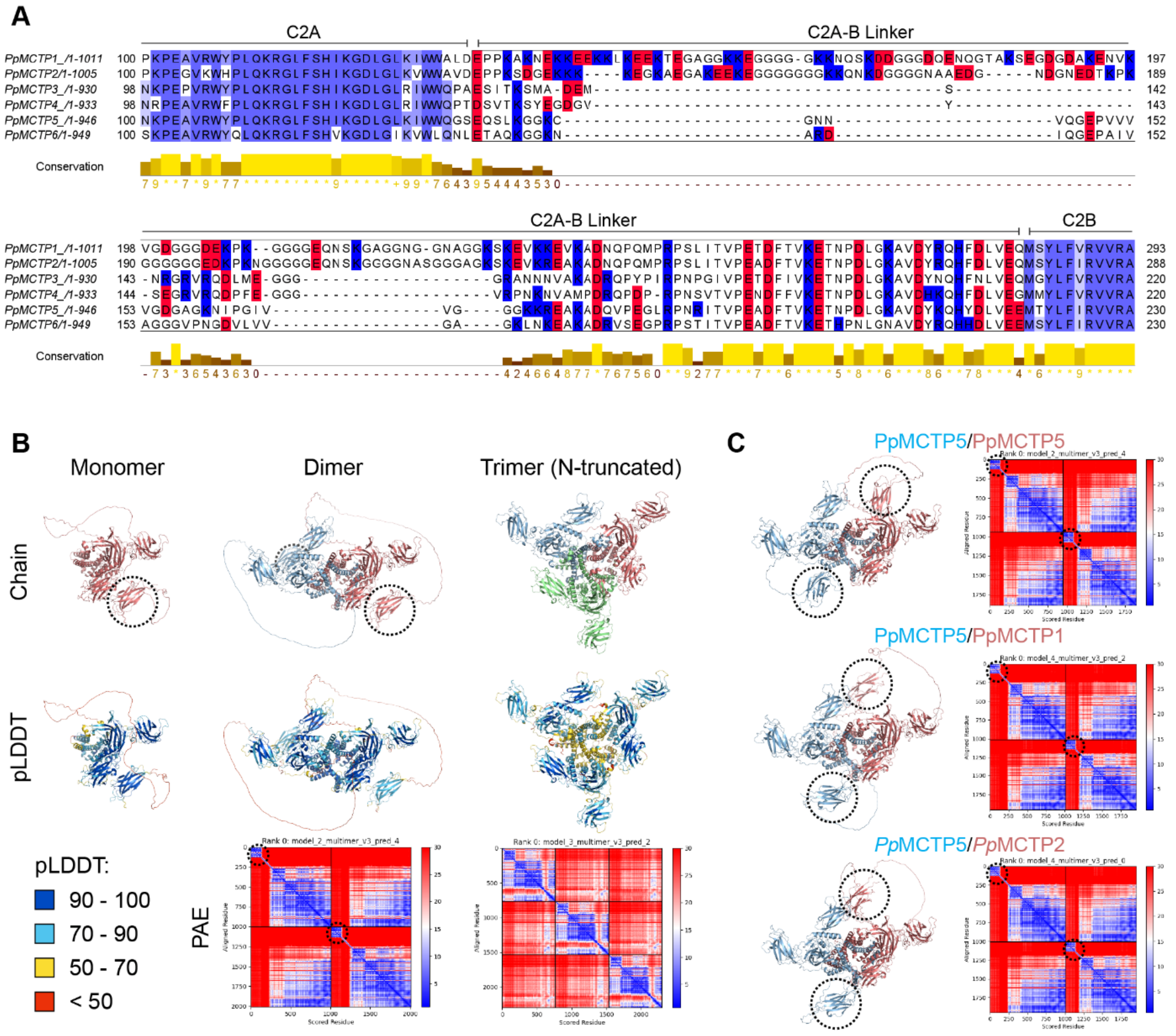


**Fig. S4. Sequence and structural analysis of *Pp*MCTPs.**
(**A**) Alignment of C2A-B linker domains. While C2 domains are highly conserved, the linker domains vary in length and residue composition. These linkers are glycine-rich and polyampholytic, with evenly distributed charges (blue: positive, red: negative). This composition suggests high solubility and flexibility, with a propensity for unspecific intra or intermolecular interactions. Identical residues in C2A and C2B are colored in blue. **(B)** AlphaFold2-predicted structures of monomeric, dimeric, and trimeric *Pp*MCTP2 with corresponding model confidence metrics. Dotted circles mark positionally flexible C2A domains. While the predicted local distance difference test scores (pLDDT) within C2A domains itself are high, indicating well-defined local structure, the predicted aligned error (PAE) between the C2A domain and the rest of the protein exceeds 30 Å, suggesting that the domain is positionally flexible while maintaining its domain integrity. In contrast, the C2A-B linker domains exhibits low pLDDT (< 50) and high PAE (> 30 Å), supporting intrinsic flexibility. Central helices and RHDs show increased pLDDT in the dimer compared to the monomer. In trimeric models, both pLDDT and PAE indicate reduced model confidence at the oligomerization interfaces, supporting the dimer model as the favorable oligomeric arrangement. For trimer predictions, the flexible N-terminus was truncated to reduce compute, as it does not participate in oligomer contacts. **(C)** Structure predictions reveal no substantial differences between homo- and hetero-*Pp*MCTP dimers. Besides the positional variability of C2A domains (dotted circles) and C2A-B linker domains, the predicted conformations are nearly identical and no apparent shift in confidence metrics is observed. Combined, sequence analysis and structure prediction suggest that both homo- and heterodimeric pairings of all six *Pp*MCTPs are biochemically, structurally and evolutionarily plausible.

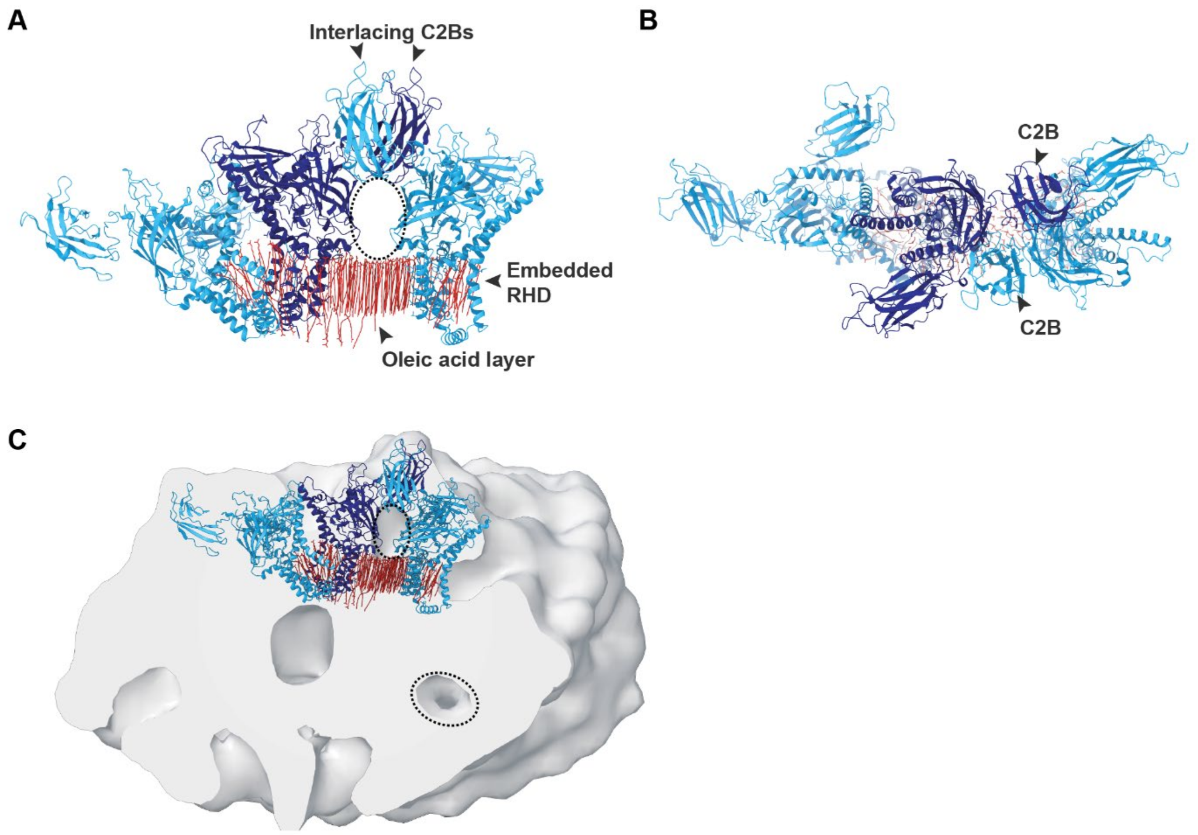


**Fig. S5. Structure prediction of trimeric *Pp*MCTPs in the presence of lipids recapitulates the experimentally determined desmotubule coating assembly.**
AlphaFold3 enables structure prediction of proteins in the presence of selected ligands (*63*). When added in excess, the fatty acid ligand oleic acid approximates membrane-like chemical properties and improves prediction accuracy for membrane proteins. (**A**) Side and (**B**) top view of predicted trimeric *Pp*MCTP2 (monomers shown in alternating light and dark blue) with 140 oleic acid molecules (red) forming a layer around the reticulon homology domains (RHDs). C2B domains interlace as observed in the experimentally resolved desmotubule-coating assembly. Dotted ellipse marks unoccupied spot between membrane and interlaced C2Bs. C2A and the flexible C2A-B linker are excluded. (**C**) Angled cross-section of the experimental map with docked trimeric AlphaFold3 model. Unoccupied spots underneath interlaced C2 domain pairs align well between model and experimental map (dotted circles).

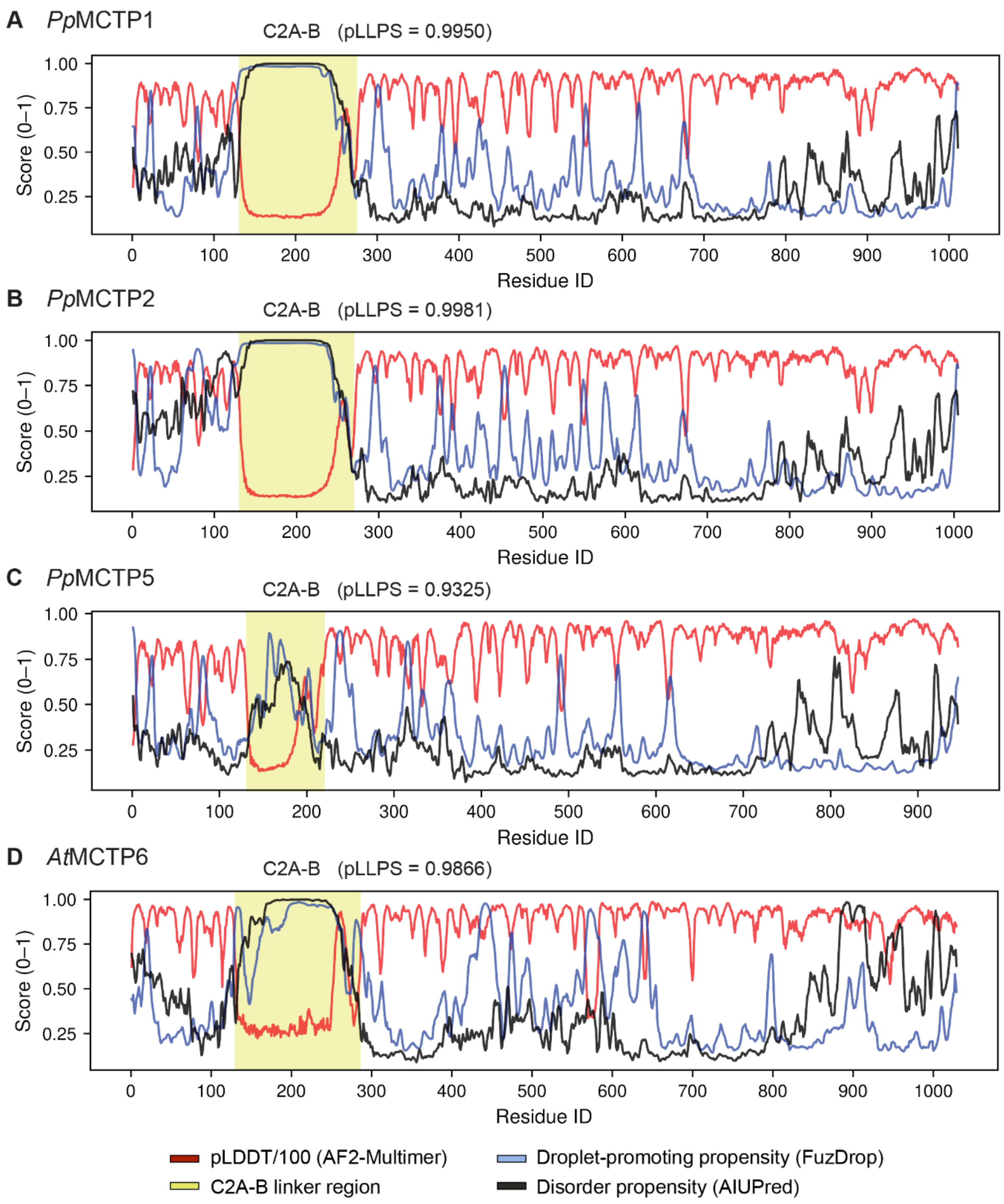


**Fig. S6. In silico profiling of sequence-encoded biophysical properties of full-length MCTPs.** (**A**) *Pp*MCTP1, (**B**) *Pp*MCTP2, (**C**) *Pp*MCTP5, and (**D**) Arabidopsis *At*MCTP6. Shared y-axis shows per-residue confidence scores from AlphaFold2-Multimer predictions (pLDDT/100, red), predicted droplet-promoting propensity (pDP) from FuzDrop (blue), and predicted disorder propensity from AIUPred (black). C2A-B linker domains are indicated in yellow. Liquid-liquid phase separation potentials (pLLPS, FuzDrop) of C2A-B linker domains are stated for each homolog.

**Table S1. Proteomics-derived list of desmotubule coating assembly candidate proteins.**
(provided as auxiliary supplementary material)

**Table S2. *Pp*MCTP gene identifiers and protein names.**
Gene identifiers, protein names, rank in proteome sorted by enrichment in PD fractions, and length of C2A-B linkers in amino acids. *Pp*MCTP1, 2, 3, 4, and 5 are found in the proteome with *Pp*MCTP1, 2 and 5 being most abundant.

| Protein name | Gene ID v6 | Gene ID v3 | Proteome rank (Gombos et al. 2023) | C2A-B linker length |
|---|---|---|---|---|
| *Pp*MCTP1 | Pp6c10_5830 | Pp3c10_11080 | 158 | 150 aa |
| *Pp*MCTP2 | Pp6c14_13840 | Pp3c14_25200 | 20 | 145 aa |
| *Pp*MCTP3 | Pp6c25_4790 | Pp3c27_540 | 319 | 80 aa |
| *Pp*MCTP4 | Pp6c16_4930 | Pp3c16_9250 | 318 | 80 aa |
| *Pp*MCTP5 | Pp6c25_4800 | Pp3c27_520 | 5 | 88 aa |
| *Pp*MCTP6 | Pp6c16_4910 | Pp3c16_9260 | Not detected | 88 aa |
